# Development of a photostable pH biosensor based on mStayGold

**DOI:** 10.64898/2026.03.06.710027

**Authors:** Michael Chang, Kei Takahashi-Yamashiro, Takuya Terai, Robert E. Campbell, Kelvin K. Tsao

## Abstract

pH-sensitive fluorescent proteins (FPs) play a crucial role in investigating pH-related cellular processes, such as endocytosis and exocytosis. Existing pH-sensitive FPs generated from *Aequorea victoria* green fluorescent protein (GFP), such as superecliptic pHluorin (SEP) and Lime, have been widely employed to study these processes, but suffer from low photostability. Here, we report the development and characteristics of serapH, a genetically encodable pH biosensor with improved photostability compared to GFP analogues, which we generated using mStayGold as a scaffold. To aid in the development of serapH, we developed a method for screening pH-sensitive FP variants by directly evaluating both brightness and pH sensitivity in bacterial colonies on agar. This significantly increased the number of colonies that could be screened per round and reduced the time needed per round. The photostability of serapH should improve spatiotemporal resolution by increasing tolerance to higher excitation intensities and longer imaging durations, thereby expanding the range of applications of pH-sensitive FPs.

## Introduction

Cellular pH is a fundamental component of many biological processes. Synaptic vesicles, for instance, have a pH of approximately 5.5 in their lumen^1^. When these vesicles undergo membrane fusion during exocytosis, exposure to the extracellular environment causes a shift to pH 7.0-7.4 for the vesicular contents. Likewise, cellular structures meant for storage or degradation such as lysosomes and endosomes have a substantially lower pH than their surrounding intracellular environment, which in turn influences the behavior of proteins and molecules within them. Researchers can obtain valuable insights into the nature of these processes by visualizing the local changes in pH that accompany them. To this end, genetically-encoded fluorescent proteins (FPs) that are particularly sensitive to pH changes in the physiologically relevant range are important tools for cellular research^2–4^. For example, these minimally-disruptive tools can enable the visualization of synaptic events in real time^1,5^, offering new insights into neurotransmission^6,7^. Beyond synaptic dynamics, pH-sensitive FPs are highly useful for studying the trafficking of membrane proteins to acidic locales, such as lysosomal or endosomal compartments^8–10^. Using the *Aequorea victoria* green fluorescent protein (GFP) as a scaffold, current FP-based pH sensors such as Lime have achieved impressive levels of performance, with high brightness and a fluorescent response up to 80-fold between pH 5.5 and 7.4^11^. One major limitation, however, is their low photostability, which is a significant issue since vesicular events occur rapidly and high frame rate imaging is required. This is typically addressed by using total internal reflection fluorescence (TIRF) imaging, which selectively excites fluorophores near the plasma membrane at low intensity, reducing both photobleaching and background signal^12^. However, TIRF is itself limited to imaging the area near the plasma membrane, making it difficult to monitor protein trafficking to intracellular structures across the whole cell^13^.

Most FPs display some intrinsic pH sensitivity due to the equilibrium between the protonated (dim with an absorbance maximum ~400 nm) and deprotonated (bright with an absorbance maximum ~480 nm) states of the chromophore. However, the p*K*_a_ of wild-type FPs is typically too low to reliably reflect changes in physiological pH, i.e. between 5.5 and 7.4^1,5^. To develop and tune pH sensitivity in a given FP, two potential strategies can be used: structure-guided rational mutation of key residues and directed evolution. Many efforts to develop pH-sensitive FPs employ a mix of both methods, with rational targeting used to create an initial prototype and directed evolution to improve the sensor’s properties beyond that. This one-two approach was used to create the first genetically encoded GFP-based pH biosensor, super-ecliptic pHluorin (SEP)^5,14^. However, directed evolution often constitutes the most time and labor-intensive span of development. One of the most established and common methods of evolution involves generating mutant plasmid libraries and evaluating them through colony-based screening of *Escherichia coli*. Since the selection of optimal variants depends on both brightness and analyte response, colonies typically need to be screened in a two-step system: an initial selection for brightness on agar plates to identify correctly-folded proteins, followed by a secondary screening where the proteins extracted from picked colonies are assayed for their response, typically in multiwell plate format^15^ (**Figure 1a, left)**. This severely constrains both the throughput and speed of each evolutionary round. To overcome this limitation, researchers have employed a variety of automation-based screening platforms, such as fluorescence-activated cell sorting (FACS)^16,17^, or direct screening of fluorescent response for analyte-dependent biosensors in rat neurons and mammalian cells using microscopy and image analysis^18–20^. However, cost and training barriers limit the accessibility of these methods, particularly in academic research settings. On top of this, mammalian cell platforms generally have limited throughput and high specificity for their targeted analyte, with no platform to date developed for pH biosensor development.

**Figure 1.**
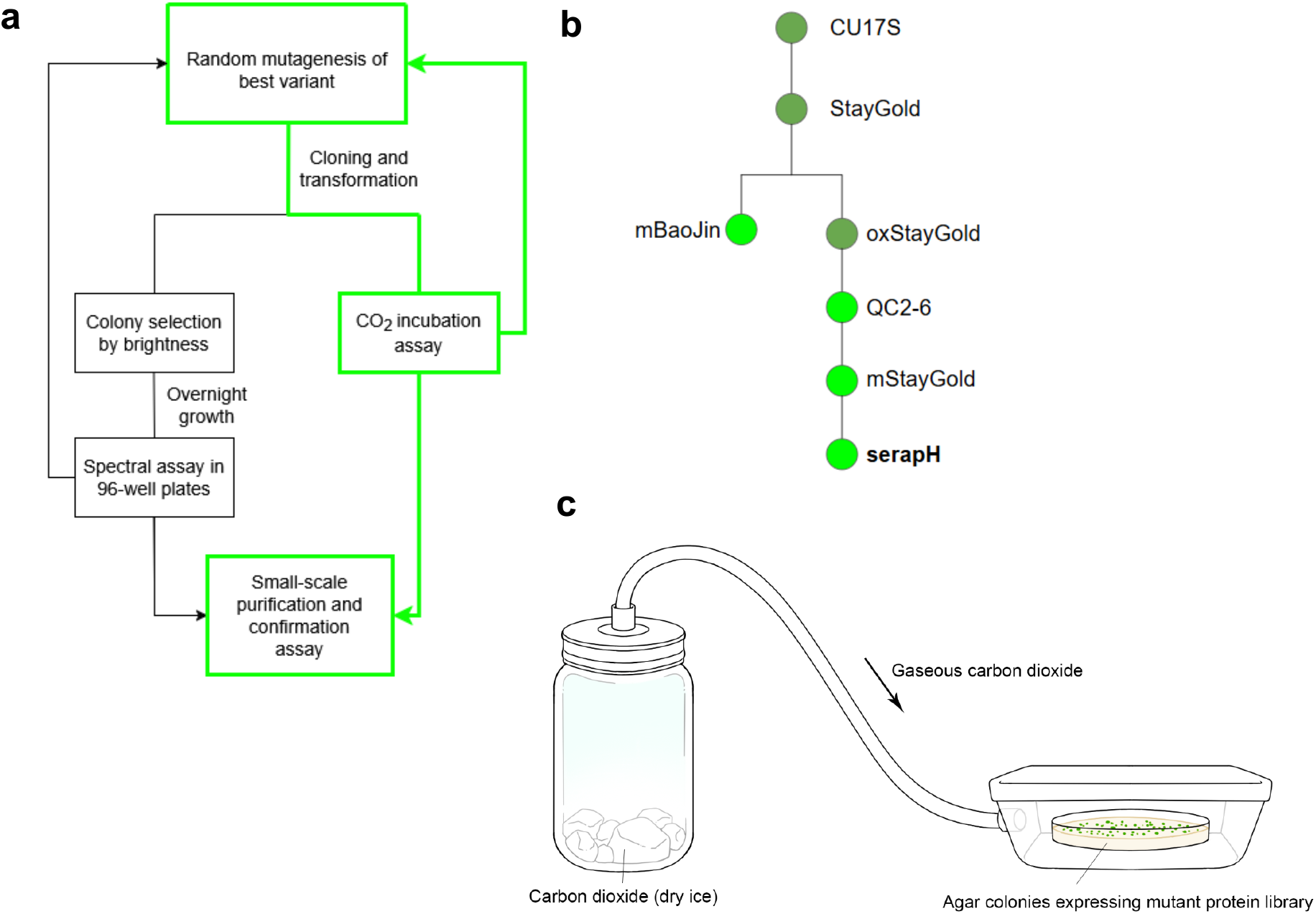
(**a**) Protein lineage of serapH development from StayGold. (**b**) Accelerated evolutionary workflow using direct screening on agar plates, showing the standard approach (left) and using the CO_2_ assay presented in this work (right, green). (**c**) Representative diagram of assay chamber for CO_2_-based screening of pH-sensitive FPs.

Here, we report the development of a new pH biosensor based on monomeric StayGold. Unlike most other green FPs, which originate from GFP first isolated from *A. victoria*, StayGold was derived from the FP CU17S originating in *C. uchidae*^21,22^, which demonstrated high photostability (**Figure 1b**). A single point mutation increased the brightness of the protein, creating StayGold, and two separate efforts have yielded the monomeric variants mStayGold(J)^22^ and mBaoJin^23^. We selected mStayGold(J) (hereafter called mStayGold or mSG) as a scaffold due to its exceptionally high photostability and suitability for protein tagging. After performing rational site-directed mutagenesis to create an initial prototype, we used directed evolution to generate a pH biosensor with a p*K*_a_ of 7.1 and approximately 21-fold intensiometric response from pH 5.5 to pH 7.4. This sensor, which we named serapH to reflect its strengths of longevity and luminance, demonstrates characteristics well-suited for live monitoring of physiological pH, including photostability similar to that of mStayGold.

## Results

### Development of an mStayGold-based pH sensor through directed evolution

Prior to performing evolution, we generated an initial serapH prototype through site-saturation mutagenesis (SSM) of two residues in the β-barrel near the chromophore using the 22C technique^24^. 22C has been shown to produce fewer redundant mutants than commonly used SSM approaches, such as NNK codons, creating a more balanced library of mutations for screening. We used a rational approach similar to that used to generate the superfolderGFP-based Lime, targeting residues 145 and 146 based on the crystal structure of StayGold^11^ (**Supplementary Figure 1**). We hypothesized that targeted mutations at these two residues would be most influential to the protein’s p*K*_a_ based on the structural characterization of sensors such as Lime^11^. To mitigate potentially low signal-to-noise ratio in the initial prototype, we screened this library through standard brightness-based colony selection, where fluorescing colonies were picked based on brightness from the agar plates and then evaluated based on their response to pH after growth in 96-well plates (**Figure 1a, left**). We subsequently identified a variant with the mutations P145E and N146A with a Δ*F*/*F* _0_ of 2.03 from pH 5.5 to pH 7.4, and termed it serapH0.1.

Next, to expedite the evolution of serapH, we developed a novel method to overcome the limitations of traditional two-step screening. In order to increase the screened library size in each round of evolution and consolidate the brightness and response-based screening steps, we designed a system that incubates bacterial colonies directly on agar in a sealed chamber with CO_2_ (**Figure 1c)**. We hypothesized that this would transiently acidify the cytosol in the bacterial colonies, highlighting the extent of pH-dependent response in the proteins expressed within them. In addition to consolidating the brightness and response screenings into a single step (**Figure 1a, right**), this method substantially increases the number of colonies that can be feasibly screened in a single round.

To test our hypothesis, we placed a Petri dish containing a pH 7.1 10 mM Tris buffer inside the incubation chamber and incubated it with CO_2_. After 15 minutes of incubation, CO_2_ had reduced the buffer pH to 4.64 ± 0.13, a result we also confirmed visually with the indicator bromocresol purple. Next, we tested pH 7.4 agar plates with colonies expressing serapH0.1, mStayGold, and Lime. We found that serapH0.1 and Lime showed substantial and reversible fluorescent responses to CO_2_ incubation, whereas mStayGold showed little to no change (**Figure 2a, b**). This response reached a maximum after approximately 15 minutes of incubation, likely corresponding to the time required for CO_2_ to reach equilibrium within the cell, and returned to baseline 15 minutes after removal from the chamber. For ease of imaging, we opted to use the restoration of fluorescence to obtain data for screening pH-dependent response. Notably, the fluorescent response of each protein on agar was substantially lower than previously reported *in vitro*, yet remained proportional to the responses of the other proteins. We also observed that, by imaging serapH with two filters at 436 nm ex./480 nm em. and 470 nm ex./510 nm em., we could quantify both fluorescence restoration in the 480 nm channel and fluorescence loss in the 510 nm channel, as a product of Tyr66 phenol-phenolate equilibrium as pH returned to 7.4. After establishing the assay’s functionality and conditions, we used it as the basis for selection in several rounds of directed evolution on serapH.

**Figure 2.**
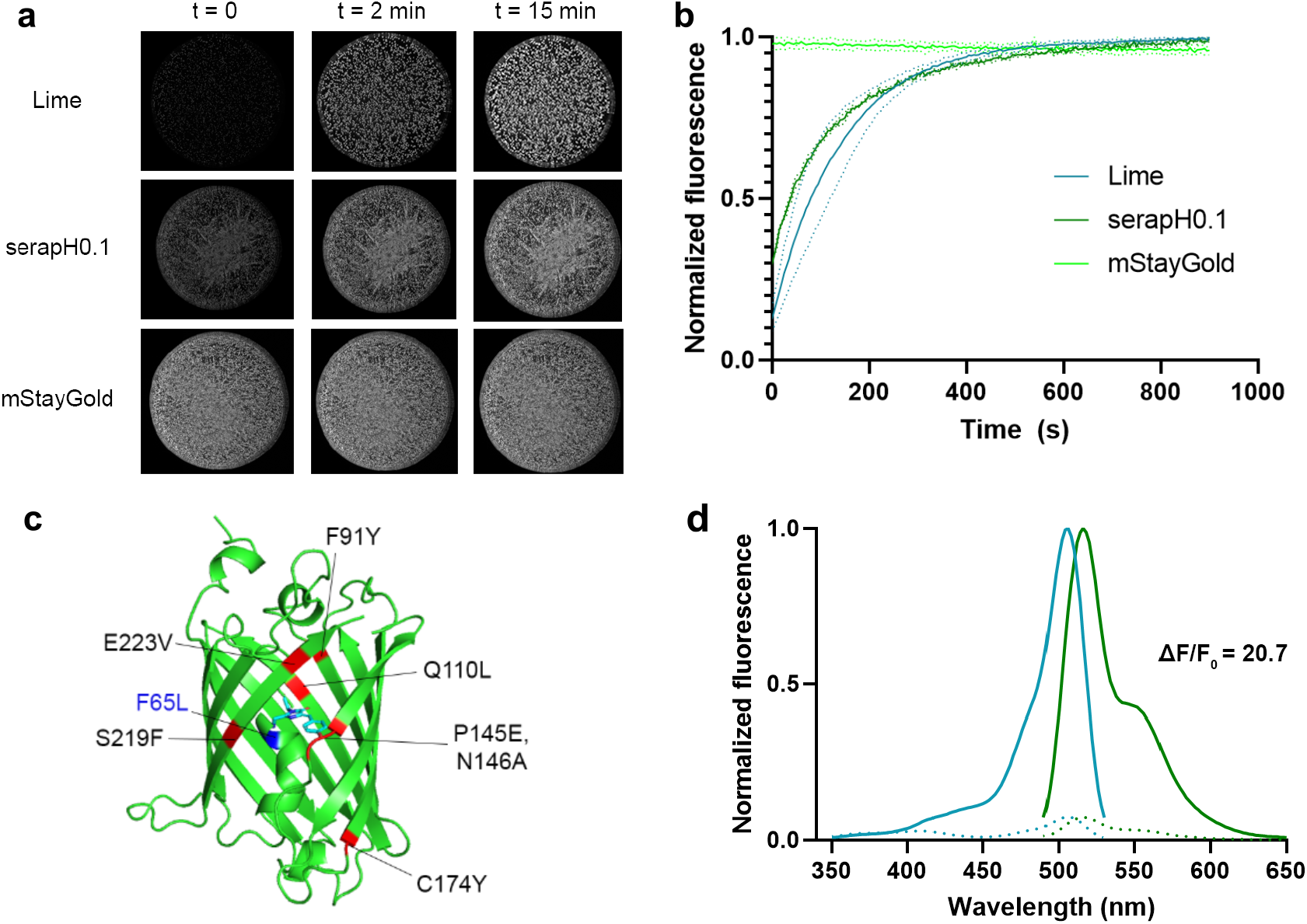
Gradual restoration of pH-dependent fluorescence in agar colonies after CO _2_ incubation at 470 nm ex/510 nm em. Imaged data from agar plates (**a**) was captured every 5 seconds, and processed in ImageJ to calculate normalized fluorescence over time (**b**) in n = 3 plates. Dotted lines represent error bars. (**c**) AlphaFold3 model^25^ of serapH1.0 with mutations marked in red, and the chromophore of StayGold (PDB ID: 8BXT) in cyan superimposed over 66Y. F65L, which was found during evolution but later reversed, is marked in blue. (**d)**Excitation and emission spectra of the final evolved variant serapH1.0 (pH 7.4 in solid lines, pH 5.5 in dotted lines).

After 10 rounds of random whole-gene mutagenesis and CO_2_-incubation-based screening, serapH’s fluorescent response between pH 5.5 and pH 7.4 improved to 21-fold. We initially generated libraries using mutagenic polymerase chain reaction (PCR) using a mix with a 0.1 mM concentration of manganese chloride, but found that among the best variants screened in the first 3-4 rounds of evolution (4 top variants were selected each round), a majority were template. We addressed this by incrementally increasing the mutation rate until we observed fluorescence in only 50% of the total library per round. After this adjustment, we saw more substantial improvements in the pH-dependent response with each round of mutation. Despite selecting for both brightness and response, we also observed a general decline in brightness from round to round. The final variant serapH0.9 contained eight mutations in total: two rational site-directed mutations used to generate the initial prototype serapH0.1, and six random mutations discovered through evolution (**Figure 2c**). In an effort to restore some of the original brightness, we finally performed a staggered extension process (StEP) to shuffle mutations between the evolved variant and the brighter serapH0.1^26^. We reasoned that some of the random mutations identified during evolution may have had a minor effect on response but a significant effect on fluorescence, and aimed to determine whether reversing these mutations in any combination would restore brightness to the protein. Screening of the shuffled library identified a final variant with the reversed mutation L65F, displaying 130% higher brightness and a response similar to that of the variant selected by evolution (**Figure 2d**). We termed this variant serapH1.0 and characterized the protein in detail.

### *In vitro* protein characterization

We assessed the spectral characteristics of serapH to determine whether it would function adequately as a tool for imaging pH changes in living cells compared with existing sensors. To characterize the protein*in vitro*, we prepared large-scale purifications of serapH, alongside Lime and mSG as references. At pH 7.4, serapH1.0 exhibited an excitation maximum (λ_ex_) at approximately 506 nm and an emission maximum (λ_em_) of 516 nm (**Figure 3a**). This was slightly red-shifted relative to the spectra of the original mSG protein, and its molecular brightness was also lower as a result of evolution. Between pH 5.5 and pH 7.4, we calculated the Δ*F/F* _*0*_ of serapH to be 20.7. Additionally, we performed a pH titration, which indicated a p*K*_*a*_ of 7.1 and a Hill coefficient n_H_ of 0.81 (**Figure 3b**). The quantum yield of serapH was determined to be unchanged between the high and low-pH states, remaining at 0.94 (**Table 1**). In turn, the loss of brightness at low pH was found to be almost entirely due to the shift in apparent extinction coefficient at 508 nm, which was determined to be 55,000 M^−1^ cm^−1^ at pH 7.4 and 6,500 M^−1^ cm^−1^ at pH 5.5. These values indicate a major shift toward the anionic chromophore species at pH 7.4, which is consistent with our characterized 7.1 p*K*_a_ value. The molecular brightness of serapH, at 52 mM^−1^ cm^−1^ (EC * QY/1000), was diminished from the original mStayGold (140 mM^−1^ cm^−1^) but remained higher than Lime (35 mM^−1^ cm^−1^). We also obtained absorbance spectra and observed two peaks at 398 and 508 nm, characteristic of the phenol and phenolate forms of Tyr66, respectively (**Figure 3c**). These characteristics showed suitability for assaying physiological pH, and to further distinguish serapH, we also addressed the key property of photostability that motivated our initial choice of mStayGold.

**Table 1.**
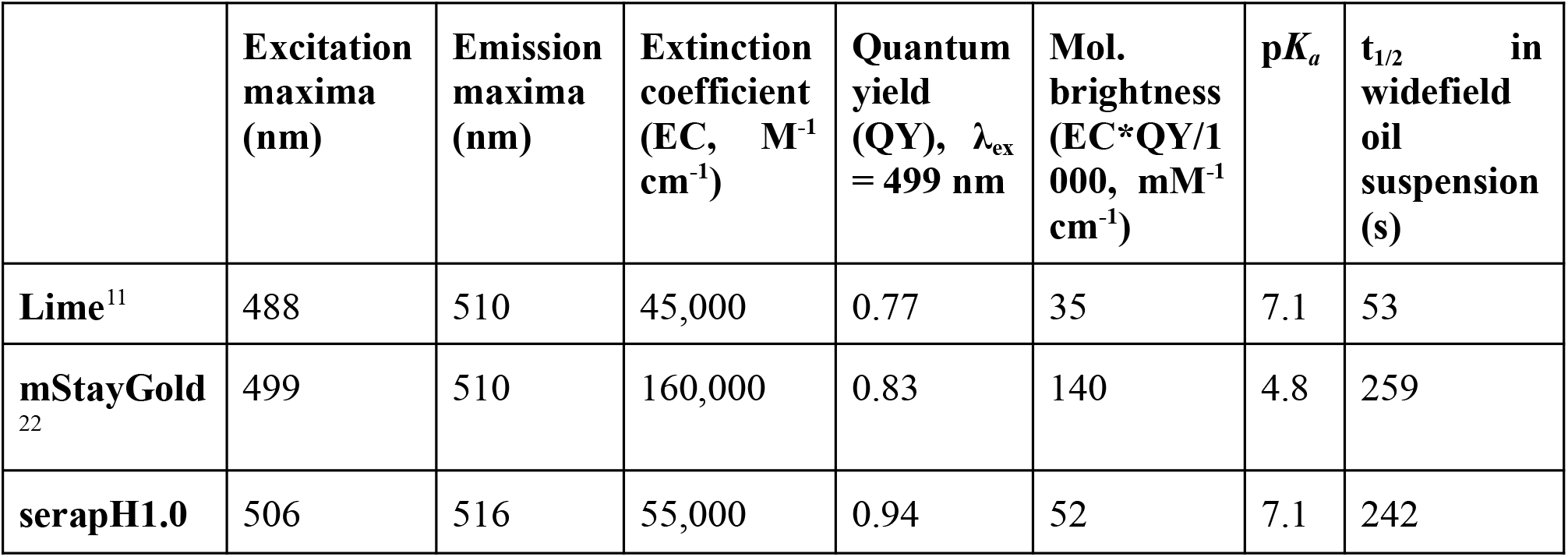
Spectral characteristics of serapH and controls. Characterized data for Lime and mStayGold were obtained from their respective publications, except for t1/2, which was measured experimentally.

**Figure 3.**
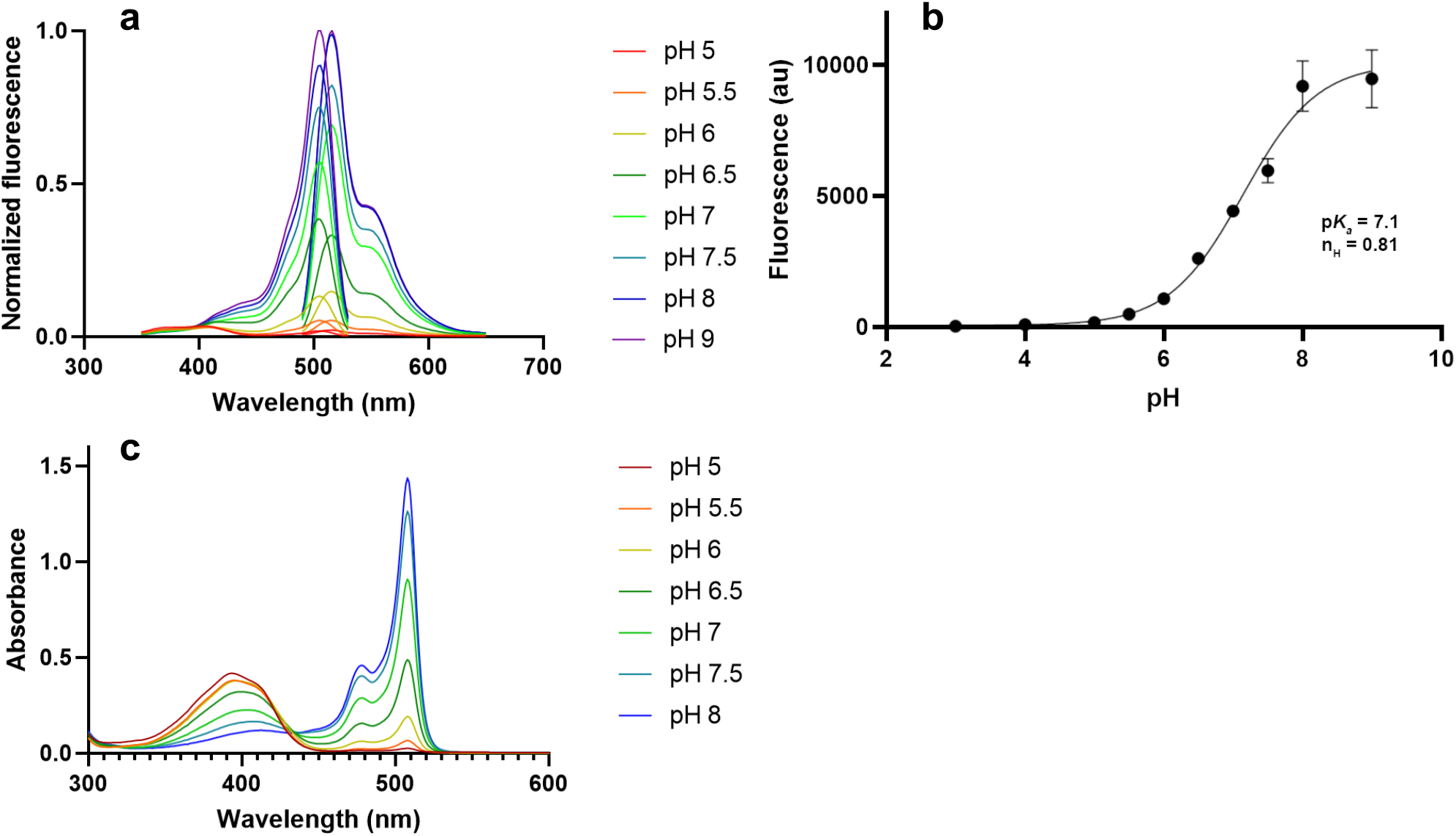
Spectral characterization of serapH1.0. (**a**) Emission and excitation spectra of serapH as a function of pH. The excitation spectra were taken by scanning from 350-530 nm with emission read at 575 nm, and the emission spectra were taken by scanning from 490-650 nm with excitation read at 455 nm. (**b**) pH titration curve of serapH1.0 using emission values averaged from 500-540 nm. p*K*_*a*_ = 7.1 and Hill coefficient n_H_ = 0.81 were calculated using a sigmoidal 4PL fit (GraphPad Prism). (**c**) Absorbance spectra at various pH values, 15 µM protein concentration.

### Characterization of photostability

Since we selected mStayGold as a scaffold for its high photostability relative to existing GFP-based sensors, we directly compared the photostability of evolved serapH with mSG and Lime to see if the photostability had been retained during directed evolution. We evaluated photostability in two ways: by illumination of purified protein solutions suspended as droplets in mineral oil, and by illumination of HeLa cells expressing the protein as a fusion to histone 2B. In both cases, protein samples were imaged continuously for 30 minutes using a widefield microscopy, with 488 nm excitation from a light-emitting diode (LED) illuminator, resulting in an irradiance of approximately 4.8 mW/cm^2^ at the plane of the microscope stage. Using measurements from imaging protein droplets in mineral oil, we observed that the photobleaching half-life (*t*_*1/2*_) of Lime was 53 seconds, while serapH and mStayGold were similar at 242 and 259 seconds, respectively (**Supplementary Figure 1**). We observed even greater photostability in the H2B fusions in mammalian cells, which exhibited only minor photobleaching over 25 minutes under our excitation conditions (**Figure 4a, b**). In both cases, serapH displayed photostability that was slightly lower than that of mStayGold, but still much higher than that of Lime.

**Figure 4.**
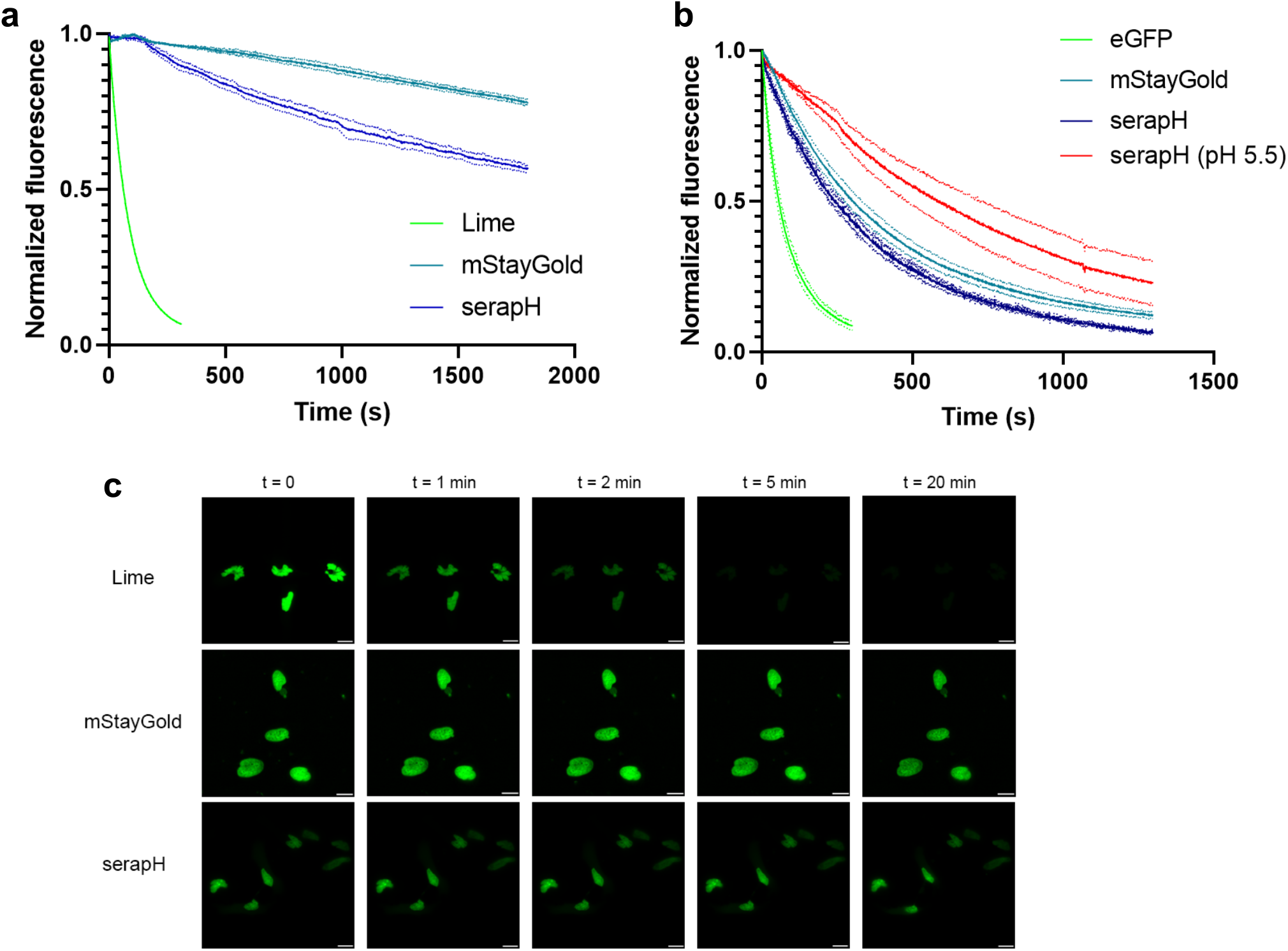
Photostability measurements of serapH and controls. Photostability was assayed through continuous illumination of the protein in a histone 2B fusion in HeLa cells (**a**) or in purified protein suspended in mineral oil (**b**) at 488 nm excitation and 518 nm emission. Curves shown represent three biological replicates (n = 3 independent experiments). (**c**) Representative time-lapse of photobleaching in H2B fusions. Scale bars represent 20 µm.

## Discussion

Here, we have reported the development of serapH1.0, a highly photostable pH sensor based on mStayGold^22^, and demonstrated a new methodology for selecting and evolving fluorescent protein-based pH sensors. SerapH1.0 has high molecular brightness, with 52 mM^−1^ cm^−1^ to 35 mM^−1^ cm^−1^ of Lime. With a p*K*_*a*_ of 7.1, it is closely aligned with existing pH sensors, such as SEP at 7.2 and Lime at 7.1. Most importantly, serapH overcomes the fundamental limitation of low photostability inherent in GFP-based sensors. Our demonstration of serapH’s photostability showed*t* _*1/2*_ = 242 seconds to bleach to half the original fluorescence in our protein droplet assay, vs 53 seconds for Lime. We observed even higher photostability in HeLa cells with H2B fusions, which may be attributed to differences in the redox state of the two environments, or additional photon absorbance from cell membranes in the H2B fusions. With its resistance to photobleaching, serapH should be able to withstand longer time-lapse image acquisition while supporting higher excitation intensities and lower gain. These qualities make it particularly well-suited for applications that demand continuous imaging at relatively low expression levels^27^. They also expand the possibilities for visualizing intracellular protein trafficking, as they enable the use of super-high-resolution microscopy techniques^21,22^ that were previously not feasible due to the high risk of photobleaching ^13^.

Although we lack a crystal structure to explain the p*K*_*a*_-altering mechanism behind the mutations we discovered over the course of developing serapH1.0, there is a plausible explanation based on the structural characterization of Lime. In Lime, the mutation S147D was found to have inverted orientation, with the polar side chain facing into the β-barrel rather than outwards ^11^. It is hypothesized that direct hydrogen bonding interactions between the chromophore and this aspartic acid residue are the primary cause for the raised p*K*_*a*_ of Lime relative to its scaffold (**Supplementary Figure 1**). We used this as the basis for our own site-saturation mutagenesis of residues 145 and 146, which led to the discovery of the P145E mutation in serapH0.1. A similar interaction between this glutamic acid residue and the mStayGold chromophore is likely present in serapH, and has also been further tuned by the additional mutations to the β-barrel found through evolution.

The CO_2_-based selection assay we have demonstrated is also an improvement to the directed evolution workflow of pH-sensitive FPs. The benefit of this system is twofold: we can directly screen proteins for both pH response and brightness in agar colonies, thereby scaling the size of the screened library while efficiently evaluating protein functionality in a single step.

Simultaneously evaluating brightness and response also allowed us to easily optimize the mutation rate during library generation. Compared to other high-throughput screening methods such as FACS, our CO_2_ incubation method is relatively low-tech and low-cost, requiring only an airtight chamber to house agar plates and a light source and camera to capture fluorescence emission from the colonies. Using this method, the time and labor required per round for a pH-sensitive FP could be reduced while increasing the likelihood of identifying positive mutations within each library.

Like other screening methods using crude lysates, the CO_2_ assay often conflates expression level with brightness when assessing colony fluorescence on agar. This is evidenced by the loss of molecular brightness in serapH (52 mM^−1^ cm^−1^) relative to mSG (140 mM^−1^ cm^−1^), despite continuous selection for both brightness and response across evolutionary rounds. We observed reduced fluorescent response for proteins in colonies compared to *in vitro*, which was consistent and proportional in both the controls and mutagenic libraries. One potential cause may have been the CO_2_’s inability to penetrate the full depth of the colony, leading to some cells being inadequately acidified. Visual artifacts in edge and low-brightness colonies also tended to overrepresent them as more responsive, which we mitigated by applying a brightness threshold in subsequent rounds. Though we were able to address these issues, mainly by employing a confirmation assay in which selected colonies are expressed at a larger scale, they limit the CO_2_ assay from fully reflecting a library’s pH sensitivity and brightness.

While its current performance is already adequate for cellular imaging, serapH still shows room for improvement in both brightness and response compared with other green-fluorescent pH biosensors. In addition to its diminished molecular brightness, serapH’s 21-fold fluorescent response remains modest in comparison to its GFP-based counterparts, which have demonstrated response levels upwards of 60-fold. The mechanistic reason for this may be reflected in the difference in the apparent Hill coefficients between these sensors, which describes the rate of fluorescence change as pH shifts near the p*K*_*a*_ threshold. The Hill coefficient of serapH is n_H_ = 0.81, which is substantially lower than the 1.9 of SEP and 1.6 of Lime. The abnormally high n_H_ values of SEP and Lime suggest that, in going from low to high pH, there may be a secondary deprotonation event that acts cooperatively with the deprotonation of the chromophore^19^. Since our prototype-generation strategy followed Lime’s rationale, which in turn used mutations from SEP, it stands to reason that serapH could also gain this cooperative interaction through further evolution and optimization. The development of other FP-based biosensors suggests that significant increases in molecular brightness without sacrificing response can be achieved through further evolution^28,29^, which would strengthen serapH in applications requiring minimally disruptive expression, such as protein tagging^8,27^. It is also important to note that while the molecular brightness of serapH is diminished relative to the original mStayGold, it is still substantially higher than Lime, meaning its brightness is not practically limiting for any existing application of Lime/SEP. To further improve the brightness and response of serapH, we plan to explore adjustments to the screening process, such as expansions to the incubation setup to incubate and image several plates in parallel. Finally, while serapH1.0 already shows some ratiometricity due to its lack of long Stokes-shift emission in the protonated state, we could use our evolution setup to create a more effective ratiometric serapH. By screening plates with an additional 436 nm/480 nm excitation filter, we could also improve the brightness of serapH’s protonated form at low pH and develop a ratiometric variant for quantitative pH measurement.

Here, we have developed serapH, a photostable pH biosensor generated from the StayGold lineage, along with a new screening system to facilitate the rapid evolution of pH-sensitive FPs. With its exceptional photostability, serapH should enable high-resolution imaging of vesicular fusion and other endocytic events over much longer time scales. Likewise, the pH-sensitive FP screening system we have demonstrated addresses some of the limitations of standard screening methods. We aim to apply the methodology developed here to further improve serapH and other relevant pH-sensitive proteins. Ultimately, we believe serapH will offer new insights into the dynamics of membrane protein trafficking to intracellular compartments, particularly through the enabling of super-resolution imaging modalities previously inaccessible due to the photostability limitations of GFP-based sensors.

## Methods

### General materials

Synthetic oligonucleotides were purchased from Thermo Fisher Scientific. The pBAD base plasmid encoding mStayGold was obtained as a gift from the Miyawaki research group. Phusion High-Fidelity DNA Polymerase (Thermo Fisher Scientific) was used for routine polymerase chain reaction (PCR) amplification and single-site-directed mutagenesis, while Taq DNA Polymerase (New England Biolabs) was used for error-prone whole gene mutagenesis. GeneJET Miniprep and Midiprep Kits were purchased from Thermo Fisher Scientific. Bacteria were transformed by electroporation (Bio-Rad, 1652100). Proteins were expressed in *E. coli* DH10B cells (Thermo Fisher Scientific) in LB media supplemented with 100μg/mL ampicillin and 0.02% L-arabinose, and extracted using bacterial protein extraction reagent (B-PER) (Thermo Fisher Scientific) unless otherwise noted. Fluorescence excitation and emission spectra were recorded on a Spark plate reader (Tecan). Data were represented and analyzed using GraphPad Prism.

### Rational engineering and directed evolution of serapH from mStayGold

To generate the initial serapH prototype, we applied 22C mutagenesis to two residues near the chromophore. This library was screened by fluorescence on agar plates (50 µg/mL ampicillin, 0.02% w/v L-arabinose), and 192 fluorescent colonies were manually selected and grown in 1 mL of Terrific Broth (TB, 50 µg/mL ampicillin, 0.02% w/v L-arabinose) in 96-deep-well dishes overnight. Cultures were centrifuged at 4,000 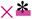 g at room temperature, then extracted with 50 uL of B-PER (Thermo Fisher Scientific). 10 µL of the B-PER extract was added to a 96-well plate and diluted with 90 µL of pH 7.4 buffer (50 mM HEPES, 50 mM MES, 200 mM NaCl) or pH 5.5 buffer (50 mM HEPES, 50 mM MES, 200 mM NaCl), after which the emission was measured using the plate reader. Excitation was measured from 350 nm to 530 nm with emission read at 575 nm, while emission was measured from 490 nm to 650 nm with excitation read at 455. A variant was selected from this screening based on fluorescent response from pH 5.5 to pH 7.4 for use as serapH0.1.

For directed evolution, the prototype was subjected to whole-gene random mutagenesis, using error-prone PCR with Taq polymerase and MnCl_2_ at a concentration tuned to generate approximately 1-2 mutations in the 800 bp gene. Blunt-end DNA fragments containing the mutant serapH gene were generated using this method. The PCR products and pBAD vector were then digested with restriction enzymes (Thermo Fisher Scientific) to yield gene fragments with sticky ends, which were subsequently ligated, cleaned, and concentrated (Zymo Research) to create a serapH library with random mutations. The libraries were transformed into DH10B *E. coli* and plated on agar plates (100 µg/mL ampicillin, 0.02% w/v L-arabinose), which were then incubated at 37 °C for 12-16 hours to allow adequate colony growth and expression.

Each plate was placed in a box and incubated with CO_2_ by feeding gaseous CO_2_ from a jar containing dry ice (**Figure 1**) for 15 minutes. After lowering the pH of the cellular environment, the plates were removed and imaged at 1 frame per 30 seconds for 15 minutes using Micro-Manager and an arc lamp (Sutter Instruments) with a 470 nm excitation/510 nm emission filter to monitor the restoration of fluorescence as CO_2_ equilibrated and the pH returned to baseline. The captured images were analyzed using ImageJ’s threshold function to create a mask highlighting each individual colony, and the region of interest (ROI) function was used to calculate their fluorescent response based on brightness values at the start and end of the elapsed time. Using these images, 4-6 colonies in each round with the highest brightness and response were selected. Selected colonies were picked and cultured in 4 mL TB (100 µg/mL ampicillin, 0.02% w/v L-arabinose), then extracted with 100 µL B-PER. 10 µL of the B-PER extract was assayed with 90 µL of each of the previous pH 5.5 and pH 7.4 buffers in 96-well optical plates (Thermo Fisher Scientific). The best-performing variant based on both brightness and response was selected and used as the template for mutagenesis in the next round.

After 10 rounds of evolution, the initial and final variants of serapH were combined to generate a library using the staggered extension process (StEP). The initial serapH0.1 prototype with two mutations and the final serapH variant evolved from it were combined into a PCR mix at 20 ng each, and a modified PCR protocol was performed with no extension step for 100 cycles^26^. PCR products were separated by agarose gel electrophoresis, and the band corresponding to the expected gene size was excised, extracted, digested, and ligated into a pBAD expression vector. This library was screened using the CO_2_ box described above to see if any combination of reverted mutations could yield a better variant. A variant was identified with a single reverse mutation that maintained a high response while restoring significant brightness, and we selected it for use as the finished sensor serapH1.0.

### Protein purification by nickel-NTA

After transforming *E. coli* DH10B with pBAD plasmids encoding the protein of interest carrying an N-terminal 6x His-tag, colonies were picked and used to inoculate a 5 mL LB (50 µg/mL ampicillin) seed culture at 37 °C and 200 rpm overnight. The starter cultures were added to 250 mL of TB (100 µg/mL ampicillin). TB cultures were shaken at 250 rpm and 37 °C until OD600 was approximately 0.6, and then induced with a final concentration of 0.02% w/v L-arabinose at room temperature and 200 rpm overnight. Cultures were pelleted at 4,000 * g, resuspended in lysis buffer (50 mM phosphate buffer, 300 mM NaCl, 1% B-PER, pH 8), and lysed by sonication with a Branson sonicator. The whole cell lysate was centrifuged at 12,000 * g and 4 °C for 30 minutes to pellet cell waste, and the supernatant was loaded onto 1 mL of nickel-NTA resin (Wako). The cell lysate was then incubated with the resin at 4 °C with rotation for 2 hours. The resin was washed with wash buffer (50 mM phosphate buffer, 300 mM NaCl, 20 mM imidazole, pH 8) for 3 × 5 resin volumes (RVs). Following this, the protein was eluted from the beads with elution buffer (50 mM phosphate buffer, 300 mM NaCl, 250 mM imidazole, pH 8), and the eluate was collected in a Falcon tube. After purification, the imidazole was removed by exchanging into storage buffer (10 mM Tris, 150 mM NaCl, pH 7.3) using a 10 kDa MWCO Amicon centrifugation filter.

### *In vitro* characterization of serapH

For characterization of excitation and emission spectra, proteins were assayed at 1 µM in pH 5.5 and pH 7.4 buffers in a 384-well optical plate (Thermo Fisher Scientific). Excitation was measured from 350 nm to 530 nm with emission read at 575 nm. Emission was measured from 490 nm to 650 nm, with excitation at 455 nm. To calculate the Δ*F/F* _*0*_, emission values between 500 nm and 540 nm (step size = 2, 20 values) were averaged from n = 3 wells for samples at pH 5.5 and pH 7.4, and calculated from the formula Δ*F/F* _*0*_ = (F_pH7.4_-F_pH5.5_)/F_pH5.5_.

For the pH titration, a series of buffers was prepared using 30 mM trisodium citrate, 30 mM sodium borate, 30 mM MOPS, and 100 mM KCl, and adjusted to pH values ranging from 3 to 11. Absorbance spectra were measured using the Shimadzu UV1800 spectrometer from 300 nm to 650 nm for proteins in the above buffer series at a protein concentration of 1 µM. To calculate molar extinction coefficients at high and low pH, the protein concentration was adjusted in 750 µL of pH 5.5 or pH 7.4 buffer until the absorbance peak at 508 nm was approximately 1. Starting from this concentration, a linear curve was measured for a series of 6 dilutions prepared at a 1:3 ratio, from which the extinction coefficient was calculated from the slope. Fluorescence quantum yield was measured with 10 µM of protein in a pH 5.5 or pH 7.4 buffer using a Hamamatsu Photonics absolute quantum yield spectrometer (C9920-02G), at an excitation wavelength of 499 nm. Quantum yield measurements were carried out in biological triplicate (n = 3), with three technical replicates per condition.

### Photostability characterization of serapH

Photostability of serapH was evaluated both in oil droplets containing purified protein and in HeLa cell nuclei expressing the protein. An IX83 wide-field fluorescence microscope (Olympus), using a pE-300 LED light source (CoolLED) and a 40× objective lens (numerical aperture (NA) = 1.3; oil), an ImagEM X2 EMCCD camera (Hamamatsu), and Cellsens software (Olympus) were used for all microscopy experiments. Imaging was conducted using 488 nm excitation/518 nm emission filters. The LED laser was measured using a Hioki 3664 optical power meter to confirm an intensity of 4.8 mW at the staging area. All photostability measurements were performed in biological triplicate.

### Purified protein droplets

Assaying of purified protein in mineral oil was done according to previously reported methods^30^. To neutralize the pH of the mineral oil for the protein suspension, a simple extraction was performed using a pH 7.3 Tris buffer. Briefly, 1 mL of mineral oil was vortexed with an equal volume of buffer, allowed to settle, and the buffer was replaced manually by pipette. This process was repeated 3-4 times until the buffer was no longer cloudy after vortexing. Purified protein was diluted in pH 7.4 or pH 5.5 buffer (5 µM for pH 7.4 and 50 µM for pH 5.5), and then suspended in a 1:100 ratio in neutralized mineral oil to a total volume of 100 µL. This mixture was vortexed at high speed for 30 seconds to disperse the aqueous protein droplets, then 10 µL was applied to a coverslip and imaged continuously at 1 frame per second for 30 minutes. Image stacks were analyzed using ImageJ to highlight individual protein droplets. Fluorescence of n = 3 individual droplets was averaged in each frame and plotted over time, and a single-decay exponential function was applied to calculate the half-life *t* _*1/2*_ for each protein.

### Cell imaging with histone 2B fusions

pcDNA plasmids containing histone H2B-serapH, Lime, and mSG were created by Gibson assembly (New England Biolabs) using the mPapaya1-H2B-6 plasmid as a backbone (Addgene #56651). HeLa (American Type Culture Collection; ATCC #CCL-2) cells were cultured in Dulbecco’s modified Eagle medium (DMEM high glucose; Nacalai Tesque) supplemented with 10% fetal bovine serum (FBS; Sigma-Aldrich) and 100μg mL-1 penicillin and streptomycin (Nacalai Tesque). Cells were seeded in 35 mm glass-bottom dishes (IWAKI). The next day, cells were transfected with pcDNA-H2B-serapH, Lime, or mSG using polyethyleneimine (PEI, Polysciences) in Opti-MEM I medium. Four hours after transfection, the Opti-MEM I solution was aspirated and replaced with 2 mL of warmed culture medium. The cells were then imaged 48-72 hours after transfection. Before imaging, the culture medium was aspirated, washed with Hank’s balanced salt solution (HBSS, Nacalai Tesque), and exchanged into HBSS. Cells expressing H2B-serapH and H2B controls were imaged continuously for 30 minutes and analyzed using ImageJ. Fluorescence of n = 3 individual nuclei was averaged in each frame and plotted over time.

## Data availability statement

The data supporting the findings of this study are available from the corresponding authors upon reasonable request.

## Author contributions

M.C. performed directed evolution and *in vitro* characterization of serapH. M.C. and K.K.T. designed and constructed the CO_2_-based screening system. M.C., K.T-Y., K.K.T., and R.E.C. designed and performed imaging-based photostability experiments. R.E.C., K.K.T., and M.C. conceived the project premise and rationale for creating an initial serapH prototype. M.C., K.K.T., T.T., and R.E.C. wrote the manuscript. K.K.T., T.T., and R.E.C. supervised research.

## Acknowledgements

This work was supported by grants from the Japan Society for the Promotion of Science (JSPS) KAKENHI (19H05633, 22H04743, 24H02267, and 24H00489 to R.E.C., 21H00273 and 23H02101 to T.T.), JST CREST (JPMJCR25T3), and the Mitsubishi Foundation. M.C. is supported by the University of Tokyo’s Global Science Graduate Course (GSGC). We thank Dr. Atsushi Miyawaki for his generous donation of the mStayGold plasmid and sequence.

