## Supplementary information for "Development of a photostable pH biosensor based on mStayGold"

### Supplementary materials

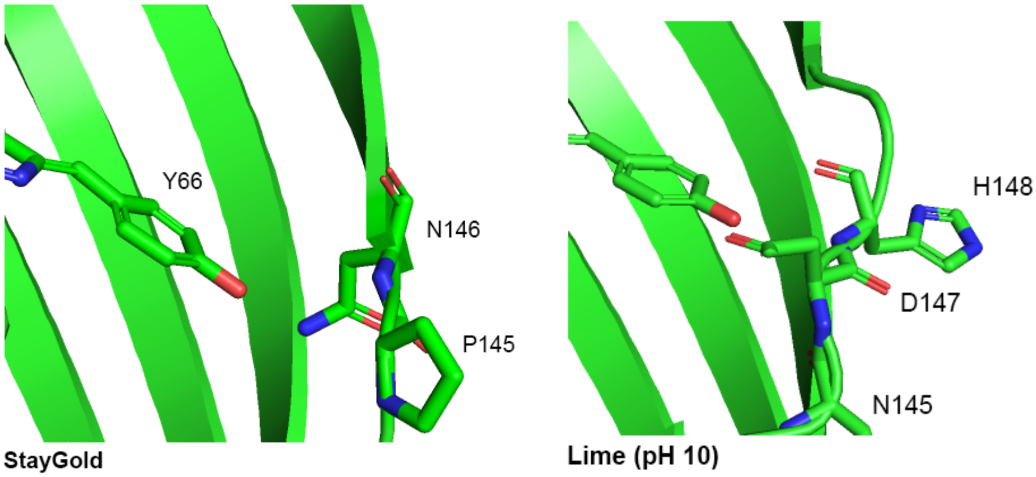


**Supplementary Figure 1.** Crystal structure comparison of the chromophore environments of StayGold (PBD ID: 8BXT, left) and Lime at pH 10 (PDB ID: 7YV3, right). The mutation D147 in Lime, whose side chain points inside the β-barrel, is thought to be a major cause of its elevated p$K_a$. We hypothesized that mutations to the analogous residues in StayGold, particularly into polar residues with negatively charged side chains, would produce a similar result.

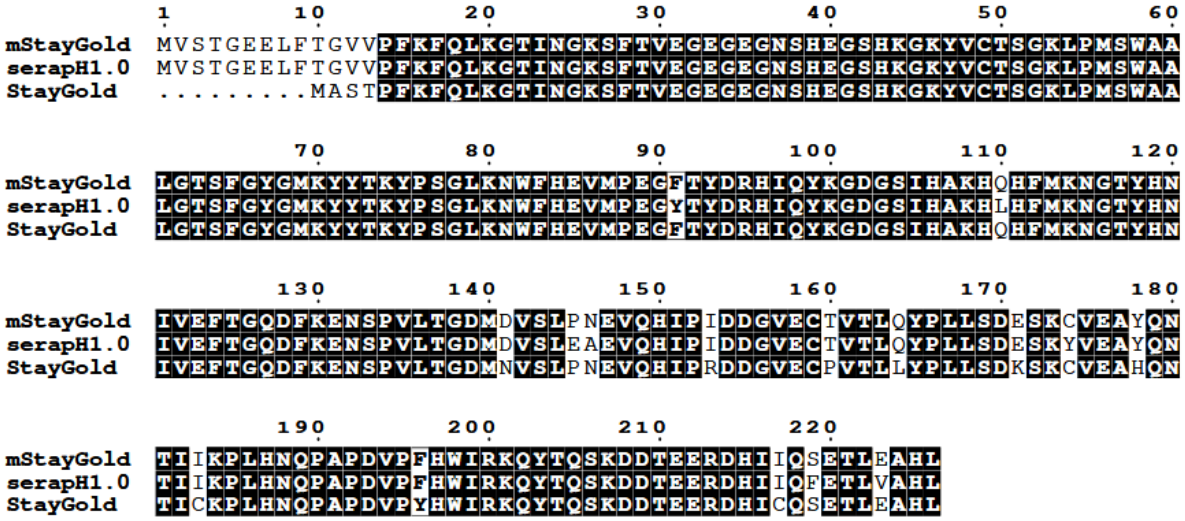


**Supplementary Figure 2.** Sequence alignment of StayGold, mStayGold(J), and serapH1.0. Alignment visualization was performed with ESPript [31].
